# Genome-wide CRISPR activation screen identifies JADE3 as an antiviral activator of NF-kB

**DOI:** 10.1101/2023.09.29.560128

**Authors:** Moiz Munir, Aaron Embry, John G. Doench, Nicholas S. Heaton, Craig B. Wilen, Robert C. Orchard

## Abstract

The innate immune system features a web of interacting pathways that require exquisite regulation. To identify novel nodes in this immune landscape we conducted a gain of function, genome-wide CRISPR activation screen with influenza A virus. We identified both appreciated and novel antiviral genes, including JADE3 a protein involved in directing the histone acetyltransferase HBO1 complex to modify chromatin and regulate transcription. JADE3 is both necessary and sufficient to restrict influenza A virus infection. Interestingly, expression of the closely related paralogues JADE1 and JADE2 are unable to restrict influenza A virus infection, suggesting a distinct function of JADE3. We identify both shared and unique transcriptional signatures between uninfected cells expressing JADE3 and JADE2. These data provide a framework for understanding the overlapping and distinct functions of the JADE family of paralogues. Specifically, we find that JADE3 expression activates the NF-kB signaling pathway, consistent with an antiviral function. Therefore, we propose JADE3, but not JADE1 or JADE2, activates an antiviral genetic program involving the NF-kB pathway to restrict influenza A virus infection.

## Introduction

The innate immune system consists of dynamic signaling networks that have substantial overlap and feedback loops to precisely tune the timing, magnitude and duration of inflammation [1]. Inflammation is a classic biological example of a double-edge sword since inflammation is critical for clearing pathogenic invaders, but uncontrolled inflammation can lead to tissue damage and is a hallmark of autoimmunity. Despite over 50 years of research, our understanding of essential regulators of innate immunity remains incomplete due to the interwoven and convoluted nature of the signaling cascades.

CRISPR/Cas9 genetic engineering has enabled scientists to perturb mammalian genomes with unprecedented precision and scale [2, 3]. Genetic screens using CRISPR/Cas9 and related technologies have increased our understanding of innate immune signaling through the systematic identification of positive and negative regulators of cytokine signaling [4–8]. While most screens have focused on loss-of-function phenotypes, modifications to the CRISPR/Cas9 system enable gain-of-function screening through the transactivation of gene expression [9]. This CRISPR activation (CRISPRa) approach enables the identification of key bottlenecks in biological systems and overcomes genetic redundancy, which may provide novel insight into the wiring of innate immune circuits.

Pathogens are excellent tools to study innate immunity since pathogens and immune systems co-evolved together. Pathogens are not only sensed by the innate immune system but also employ counter measures to circumvent cellular restriction factors. Thus, leveraging the selective pressure of pathogens such as viruses is a powerful screening approach to identify important genes involved with the immune system. Influenza A virus (IAV) is a segmented, negative sense RNA virus that is a significant cause of human disease annually with pandemic potential. While innate immune responses to IAV are critical for establishing control of viral replication, our understanding of what controls IAV infection outcomes at the individual cellular level is still not fully understood [10].

To address this knowledge gap we conducted a genome-wide CRISPRa screen to define cell intrinsic factors that can antagonize IAV infection. Our screens and subsequent validation identify both appreciated and novel regulators of IAV infection. We further demonstrate that Jade family PHD zinc finger 3 (JADE3), also called PHF16, but not the closely related paralogues JADE1 and JADE2 induces the activation of NFκB and blocks IAV replication. Overall, our results provide new insight into both host-viral conflicts and the function of poorly described host genes that may have implications beyond host-cellular defense.

## Results

### Genome wide CRISPR activation screen identifies known and novel regulators of influenza A virus infection

To identify human genes with antiviral potential against IAV, we took a gain of function CRISPR activation approach similar to our previous screens with murine norovirus [11]. Briefly, we transduced HeLa dCas9-VP64 expressing cells with the Calabrese CRISPRa library [12]. These CRISPRa pool of HeLa cells were infected with the H1N1 IAV strain Puerto Rico/8/1934 (PR8) at an MOI 1.0 in serum free media in the presence of TPCK-Trypsin to allow for viral spread (**Figure 1A**). As a control, cellular pools containing the Calabrese library were mock infected. After three days, IAV infected cultures were washed, and media replaced with serum containing media for recovery. Following a two-day recovery, cells were again infected with IAV for three days and the surviving cells were collected. sgRNAs from resistant cells were compared to mock-infected controls using STARS [13]. We consider a gene a hit if the FDR was less than 0.25. 22 genes met this criterion (**Figure 1B and Tables S1 and S2**).

**Figure 1:**
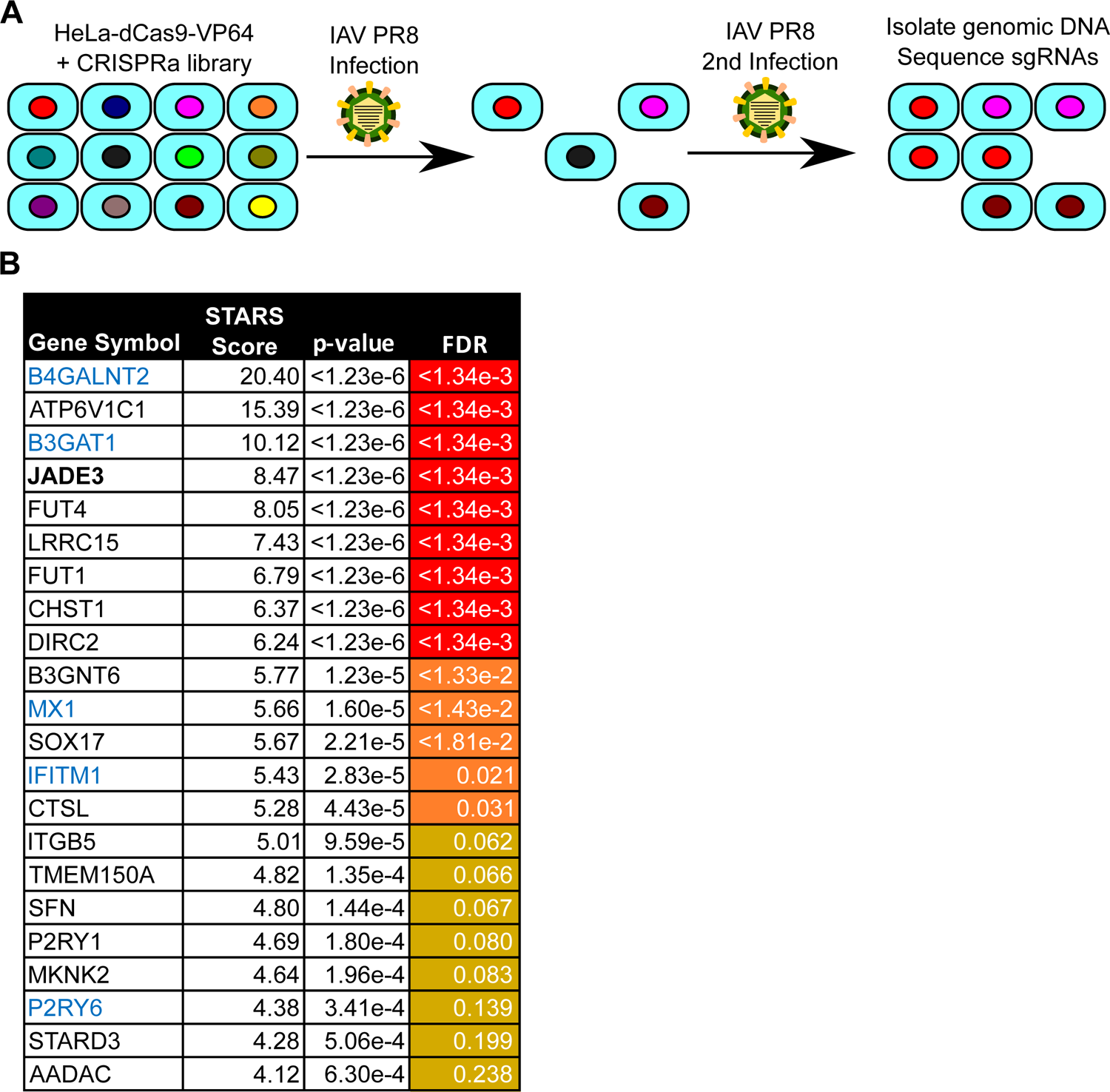
Genome-wide CRISPR activation screen identifies antiviral proteins against influenza A virus. **A.** Schematic of genome wide CRISPRa screening strategy to identify antiviral genes for influenza a virus (IAV). Sequences of the sgRNAs from cells surviving IAV challenge were quantified and compared to the relative abundance of a mock-infected sample. **B.** Table of significantly enriched genes. STARS algorithm was used to calculate STARS score, p-value, and False Discovery Rate (FDR). Genes colored in blue have previously been shown to possess anti-IAV activity. FDR scores are shown and color coded for <0.01 (yellow), <0.05 (orange), and <0.25 (gold). JADE3 (bolded) was chosen for additional validation.

Upon examination of enriched genes, we identify known antiviral genes towards IAV, including the interferon stimulated genes (ISGs) *MX1*, *IFITM1*, and *P2RY6* [14–16]. *B3GAT1* also was significantly enriched and was recently shown to broadly restrict influenza viruses by decreasing sialic acid levels at the cell surface [17]. Other hits include genes encoding the glycan modifying fructosyltransferases *FUT1* and *FUT4*, and a component of the V-ATPase *ATP6V1C1* involved in endosomal acidification suggesting possible roles for these genes inhibiting viral entry [18–20]. Finally, poorly studied genes with no known antiviral role in influenza A infection also showed significant enrichment including *JADE3* and *LRRC15*. Of these *LRRC15* has been shown to inhibit SARS-CoV2 infection and binds spike protein [21–23]. When we compare our dataset with a previously conducted CRSIRPa IAV screen, we detect only minimal overlap outside of *B4GALNT2*, the most enriched gene in each screen (**Figure S1**) [24]. This variation in data is likely due to numerous technical differences in the design of the CRISRPa screens. Overall, our screen identifies many pathways and genes appreciated to be important for IAV infection while also suggesting new genes with unknown links to IAV biology.

To validate our screening data, we individually introduced two sgRNAs per gene into HeLa-dCas9-VP64 cells targeting *B4GALNT2*, *ATP6V1C1*, *B3GAT1*, *JADE3*, *FUT4*, *LRRC15*, *DIRC2*, *TMEM150A*, *P2RY1*, and *P2RY6* and compared them to cells transduced with either an empty sgRNA plasmid or an sgRNA targeting *CD4*. We confirmed that each guide increased gene expression of the respective target gene (**Figure 2A**). We next challenged these cell lines with an IAV PR8 strain expressing mNeon as a reporter [25]. Infection was monitored for 48 hours by quantifying mNeon expression in each well every 3 hours (**Figure 2B**). Of the 10 genes tested, all 10 showed antiviral activity with at least one of the two guides and 8 genes had both sgRNAs display significant antiviral activity towards mNeon reporter IAV at 24 hours post-infection (**Figures 2C and 2D**). The presence of well-known antiviral genes amongst our enriched genes and the independent validation provides confidence that our screening dataset is robust.

**Figure 2:**
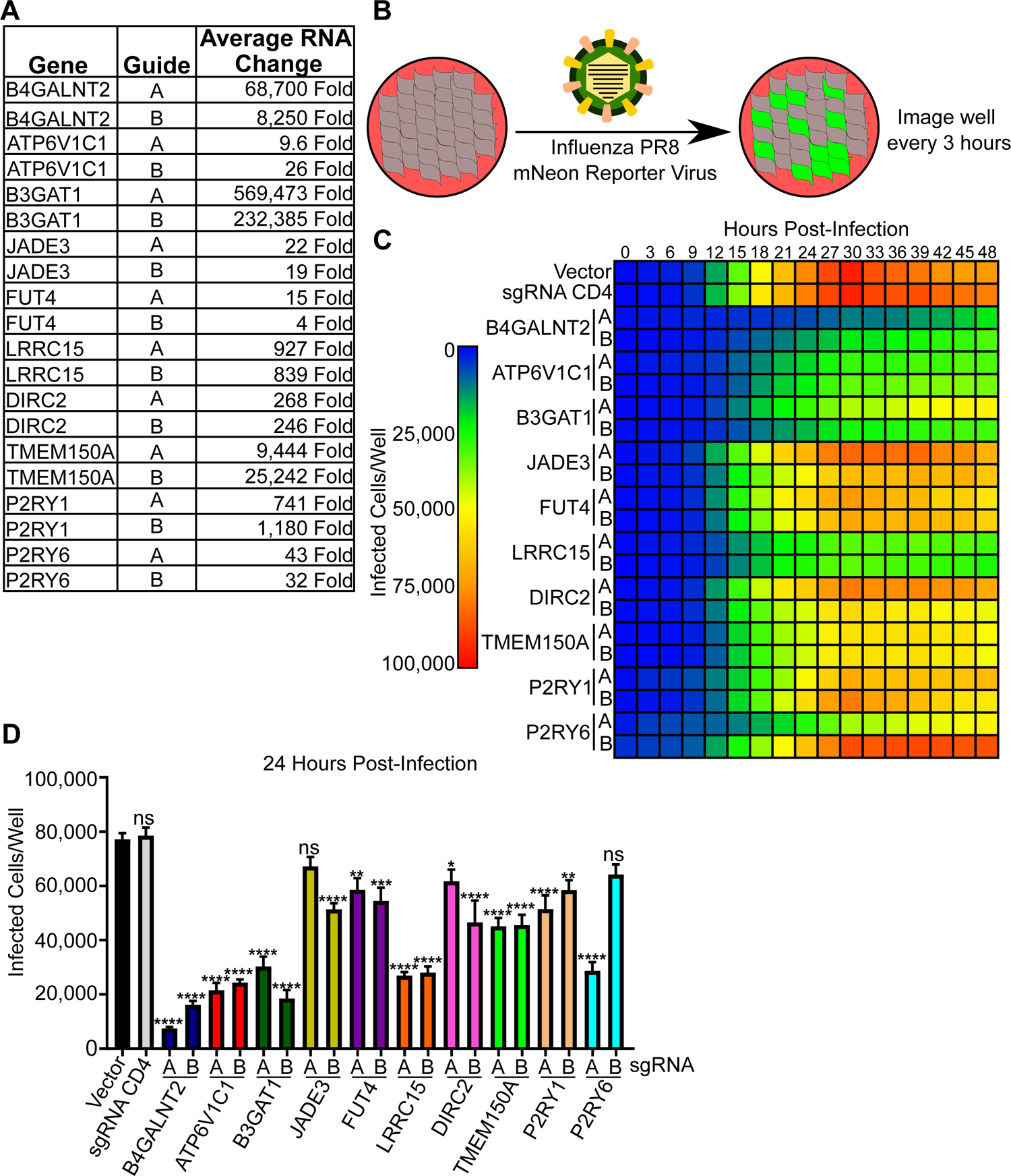
Validation of CRISPR activation screen. **A.** Table showing mRNA fold change of indicated gene in HeLa-dCas9 VP64 cells expressing guides A or B to the indicated gene. Data is shown as the average of three independent experiments. **B.** Cartoon schematic of the validation assay in which HeLa-dCas9 VP64 cells were infected with PR8 mNeon reporter virus and mNeon fluorescence was monitored every 3 hours via IncuCyte. **C.** Heatmap displaying the number of infected cells per well over a 48 hour time course for each cell line expressing the indicated sgRNA construct.. Data is the average from three independent experiments. **D.** Number of cells infected with influenza PR8 mNeon at 24 hours post infection from (**C**). The data are shown as means ± the SEM from three independent experiments, and data were analyzed for statistical differences for each cell line compared to the vector control by one-way ANOVA with Tukey’s multiple-comparison test. *, p<0.05; ** p<0.01; ***, p<0.001; ****, p<0.0001; ns, not significant

### JADE3 is sufficient and necessary to inhibit influenza A infection

Of the enriched genes with unappreciated antiviral function, *JADE3* also demonstrated antiviral activity in a recent CRISPRa screen for SARS-CoV-2 in Calu-3 cells and murine norovirus in HeLa cells [11, 26]. Together these findings suggest the anti-viral activity of JADE3 is not cell-type or virus specific. *JADE3* is a poorly studied member of the Jade family of proteins consisting of JADE1, JADE2, and JADE3. Jade family members are believed to have mostly redundant roles in the histone acetyltransferase (HAT) HBO1 complex and have been predominantly studied for their role in cancer and development [27, 28]. More specifically, the Jade family proteins direct the complex to histone H4 acetylation, which regulates gene transcription [29, 30]. Phylogenetic analysis shows distinct orthologs for all three JADE proteins extending back to jawed vertebrates (**Figure 3A**). However, while two JADE-like proteins exist in the jawless vertebrate lamprey, they do not cluster with any of the three JADE proteins, suggesting significant amino-acid replacements during the jawed-jawless divide in vertebrates. Since this divide coincides with the expansion of the immune system, we hypothesized JADE3 may have a role in the immune system [31]. Because of its novelty, its potential broad viral inhibition and its evolutionary history we selected *JADE3* for further study.

**Figure 3:**
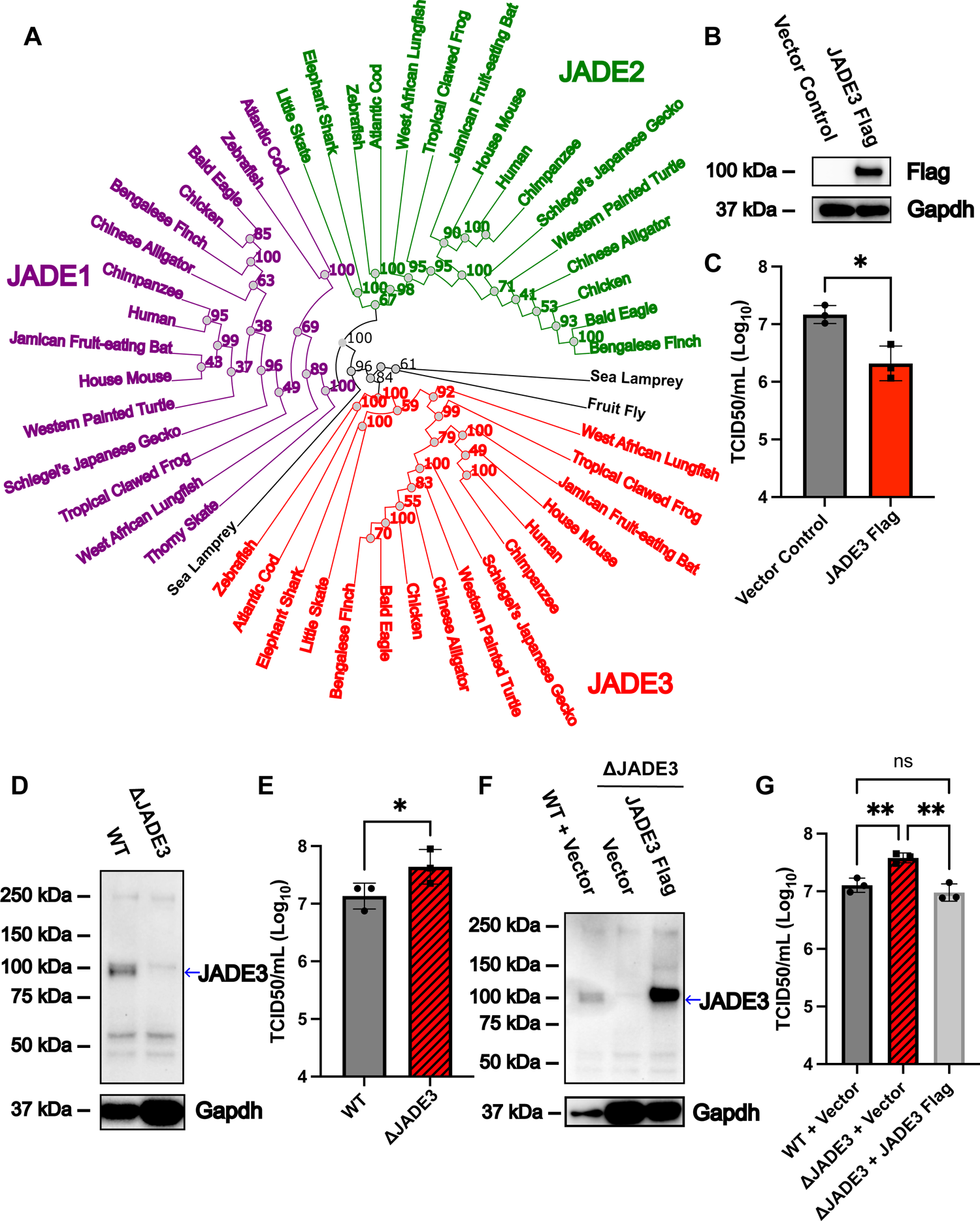
JADE3 is sufficient and necessary to restrict influenza A infection. **A.** Maximum likelihood phylogenetic tree of JADE3 orthologs. 100 bootstrap replicates were performed and bootstrap values are indicated. Branch length represents amino acid changes per site. GenBank common names are used for species. Ortholog gene groups for JADE3 (red), JADE1 (purple), and JADE2 (green) are colored. Potential JADE orthologs that do not branch with one of the three JADE proteins are in black. **B.** Representative western blot from three experiments of JADE3 Flag expression from control and JADE3 Flag transduced A549 cells. **C.** Vector control or JADE3 Flag A549 cells were challenged with WT PR8 at a MOI of 0.01. Infectious virus titer was determined by TCID50 24 hours post infection. Data was analyzed using a paired two-tail t-test. **D.** Representative western blot from three experiments for JADE3 expression in indicated cell lines. Blue arrow indicates expected molecular weight of JADE3. **E.** A549 WT or A549ΔJADE3 cells were challenged with WT PR8 at a MOI of 0.01. Infectious virus titer was determined by TCID50 24 hours post infection. Data was analyzed using a paired two-tail t-test. **F.** Representative western blot from three experiments for JADE3 expression in indicated cell lines. Blue arrow indicates expected molecular weight of JADE3. **G.** A549 control or A549ΔJADE3 cells complemented with either a vector control or Jade3 were challenged with WT PR8 at a MOI of 0.01. Infectious virus titer was determined by TCID50 24 hours post infection. Data was analyzed using a one-way ANOVA with Tukey’s multiple comparisons test. All data shown is mean +/- SD from three independent experiments. *, p < 0.05; **, p < 0.01; ns, not significant

To determine whether *JADE3* is able to inhibit a non-reporter IAV infection we generated A549 cells stably overexpressing a *JADE3* cDNA with a C-terminal 3xFLAG epitope (**Figure 3B**). A549 cells are a lung epithelial carcinoma cell line commonly used to study IAV infection [32]. A549-JADE3 cells produced less virus after 24 hours compared to A549-vector control cells. (**Figure 3C**). Next, we sought to determine the necessity of JADE3 for viral inhibition. Using CRISPR-Cas9 we generated single cell clones with indels in exon 5 of *JADE3* leading to premature stop codons (**Figures 3D and S2**). A549ΔJADE3 cells had a modest increase in viral titers 24 hours post-infection with IAV. (**Figure 3E**). Complementation with JADE3 cDNA restored IAV titers to levels comparable with the parental cell line, confirming the on-target activity of the sgRNAs (**Figure 3F and 3G**). Taken together these data indicate that JADE3 is both necessary and sufficient to optimally inhibit IAV.

### The Jade family members JADE1 and JADE2 are not sufficient to inhibit influenza A infection

JADE3 is an 823 amino acid long protein that contains 2 mid-molecule plant homeodomains (PHD, **Figure 4A**). These zinc finger domains have the capacity for the binding and recognition of histone modifications [33]. Deletion of either PHD domains, or in combination, abrogated the antiviral activity of JADE3 (**Figure 4B**). The paralogs JADE2 and JADE1 share significant sequence homology with both of the PHD domains of JADE3 (**Figure S3**). However, unlike JADE3, expression of JADE1 or JADE2 is insufficient to reduce IAV titers (**Figure 4C and 4D**). While, all Jade family members direct histone acetylation via the HBO1 complex thus regulating gene expression, our data suggests non-redundant functions of Jade family members [29, 35, 36].

**Figure 4:**
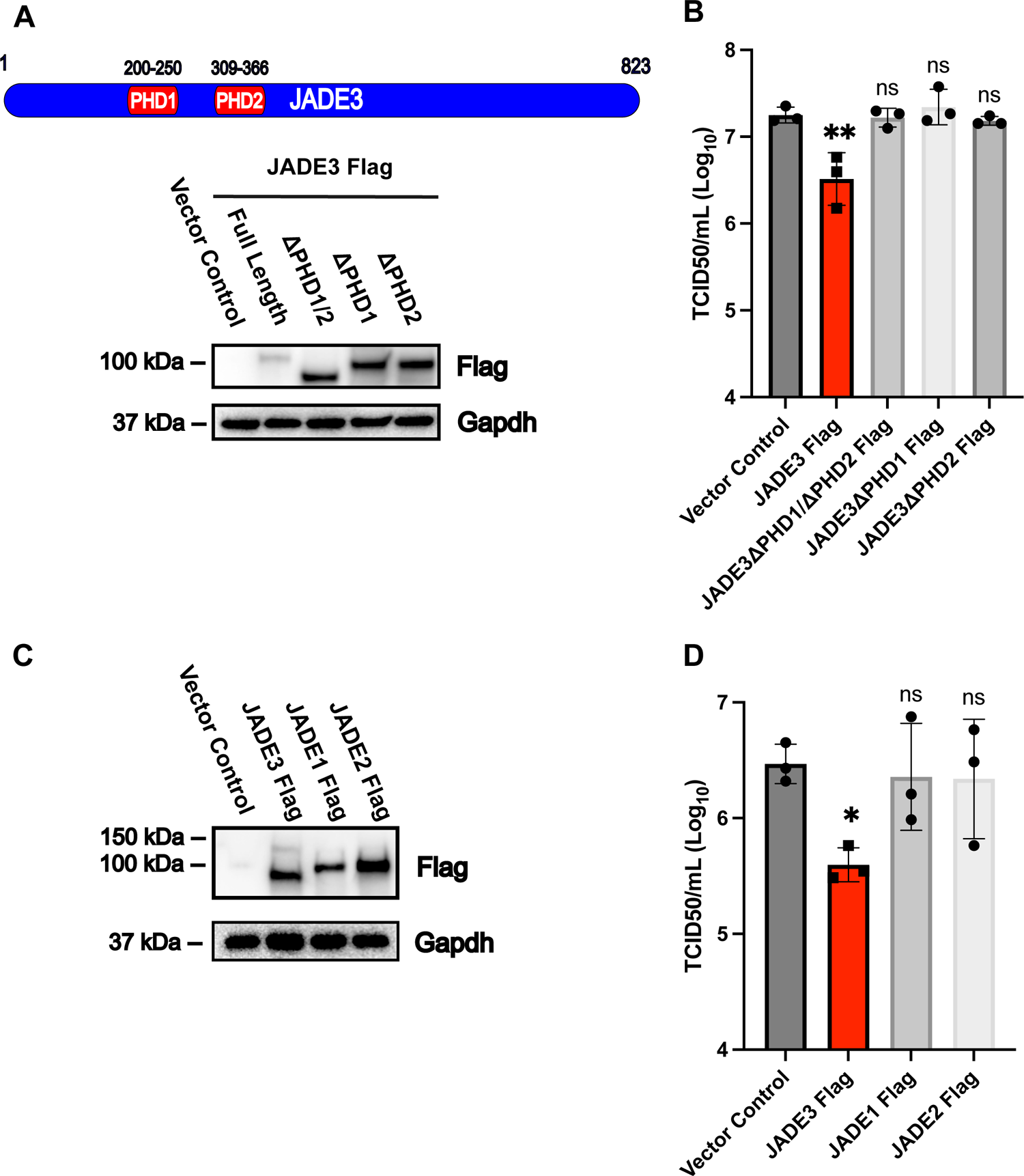
Analysis of JADE3 domains and paralogs antiviral function. **A.** Schematic of JADE3 and its PHDs. Representative western blot from three experiments of A549 cells transduced with a vector control, JADE3 Flag or JADE3 lacking both or either PHDs. **B.** A549 cells expressing either a vector control or the indicated PHD truncations were challenged with WT PR8 at a MOI of 0.01. Infectious virus titer was determined by TCID50 24 hours post infection. **C.** Representative western blot from three experiments of expression of flag tagged JADE1, JADE2, JADE3 in A549 cells. **D.** A549 cells expressing different Jade family proteins were challenged with WT PR8 at a MOI of 0.01. Infectious virus titer was determined by TCID50 24 hours post infection. All data shown is mean +/- SD from three independent experiments. Data was analyzed using a one-way ANOVA with Dunnett’s multiple comparisons test against the vector control. *, p < 0.05; **, p < 0.01; ns, not significant

### JADE3 activates expression of a TNF genetic program

The functional differences between Jade family members has greatly been understudied. Thus, we set out to define different gene expression patterns in A549 cells expressing either a vector control, JADE3, or JADE2. Principal component analysis demonstrates significant gene expression changes when either JADE3 or JADE2 is overexpressed (**Figure 5A**). Importantly, JADE3 and JADE2 samples form distinct clusters, demonstrating unique expression patterns. We hypothesized that JADE3 induces an antiviral gene expression profile distinct from JADE2. To this end we compared the differentially expressed genes from both JADE2 and JADE3 relative to vector control cells. JADE3 overexpression caused significant (FDR adjusted p-value < 0.01) upregulation of 1155 genes and down regulation of 1124 genes as compared to a vector control (**Figure 5B and Tables S3A and S3D**). JADE2 overexpression caused significant upregulation of 829 genes and down regulation of 816 genes as compared to a vector control (Table S3B). Furthermore, of the upregulated genes 716 genes were uniquely upregulated in JADE3 expressing cells and not in JADE2 expressing cells suggesting differing roles for the two JADE proteins (**Figure 5C**). A gene set enrichment analysis of the 500 most significantly increased genes in JADE3 expressing cells showed significant upregulation of the TNF/NF-kB pathway as well as other inflammatory pathways such as the IL-2 pathway (**Figures 5D, S4, S5 and Table S3C**). Since the TNF/NF-kB pathway was one of the most enriched and is well studied for its role in inflammation and viral infection we decided to explore the pathway further.

**Figure 5:**
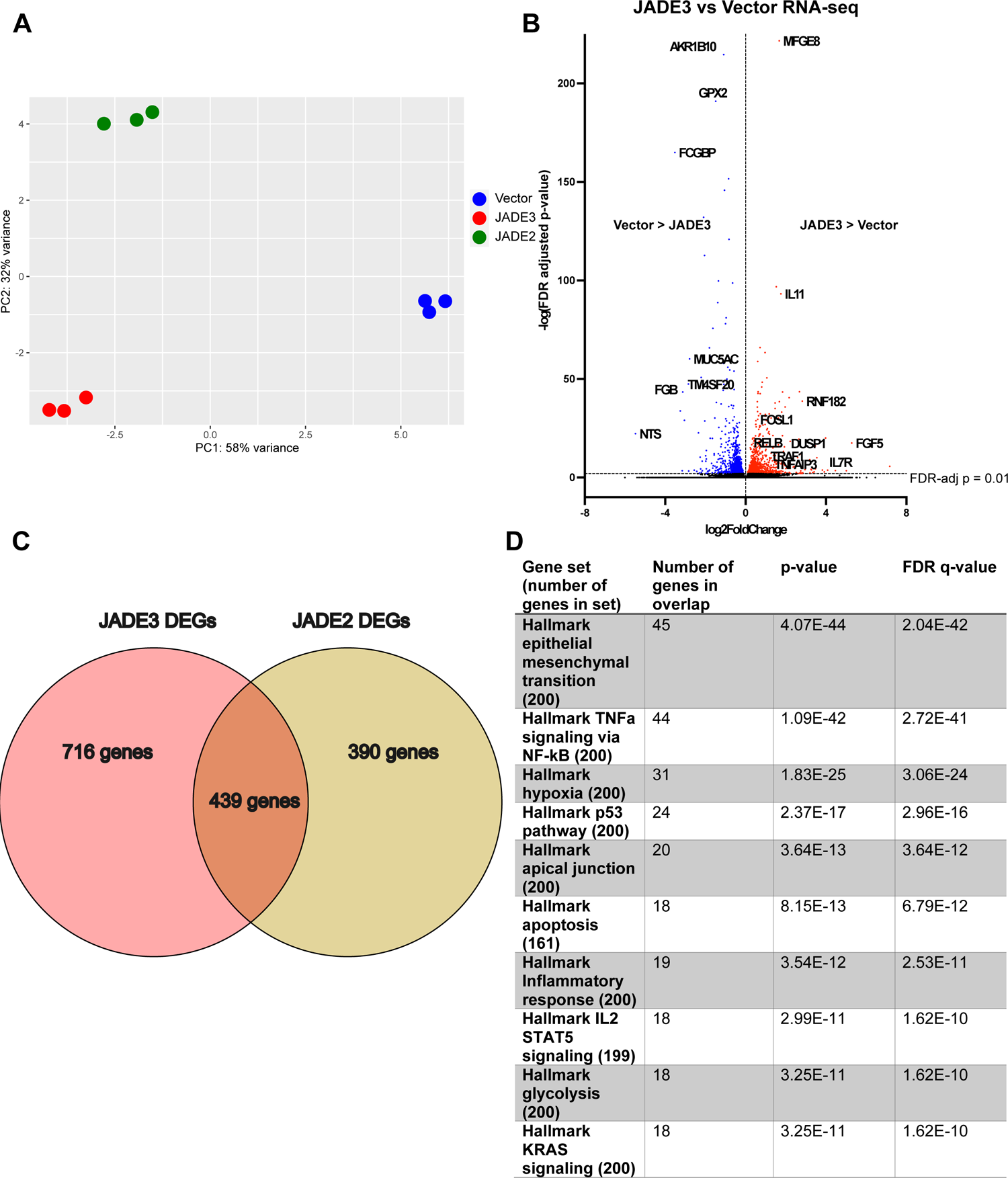
Transcriptional profiling reveals that JADE3 activates the TNF/NF-kB signaling pathway. **A.** Principal component analysis scatter plot showing differential gene expression signatures of control, JADE3 or JADE2 expressing cells. PC1 and PC2 corresponding to the principal component 1 and principal component 2. **B.** Volcano plot of RNA-seq of A549 JADE3 Flag cells vs vector control cells. An FDR-adjusted p-value < 0.01 (dotted line) was considered significant. Data is from three biological replicates. **C.** Venn diagram of differentially expressed genes (DEGs) significantly upregulated by JADE3 Flag vs vector control and JADE2 Flag vs vector control. **D.** Gene set enrichment analysis of the 500 most significantly upregulated genes in A549 cells expressing either JADE3 Flag or a vector control. The 10 most significantly enriched gene sets are shown

### JADE3 expression increases NF-kB translocation to the nucleus and phosphorylation

Focusing on the TNF/NF-kB pathway in JADE3 expressing cells, we found upregulation of canonical genes in the pathway including *RELB*, *TRAF1*, *FOSL1*, *IL6* (**Figure 6A**). In contrast to JADE3, JADE2 has only modest changes in the expression profile of the TNF/NF-kB pathway. Upon activation, NF-kB translocates to the nucleus where it activates transcription. Furthermore, NF-kB (p65) can be phosphorylated at Serine 536 which enhances its transcriptional activity and is used as a marker for active NF-kB [37, 38]. Nuclear extracts from JADE3 expressing cells had significant NF-kB p65 protein levels while it was largely absent from control cells (**Figure 6B**). JADE3 expressing cells had an increase in Serine 536 phosphorylated p65 even after treatment with recombinant human TNF-α (**Figure 6C**). Consistent with the antiviral activity of JADE3, the increase in phosphorylated p65 by JADE3 requires both PHD domains (**Figure 6D**). Importantly, expression of the other JADE paralogues is insufficient, with JADE1 expressing cells having no increase in phosphorylated p65 while JADE2 expressing cells have a modest increase in phosphorylated p65, consistent with a moderate increase in some NF-kB pathway genes (**Figure 6E**). Taken together, our data demonstrates that expression of JADE3 leads to activation of the NF-kB pathway that likely results in an antiviral state.

**Figure 6:**
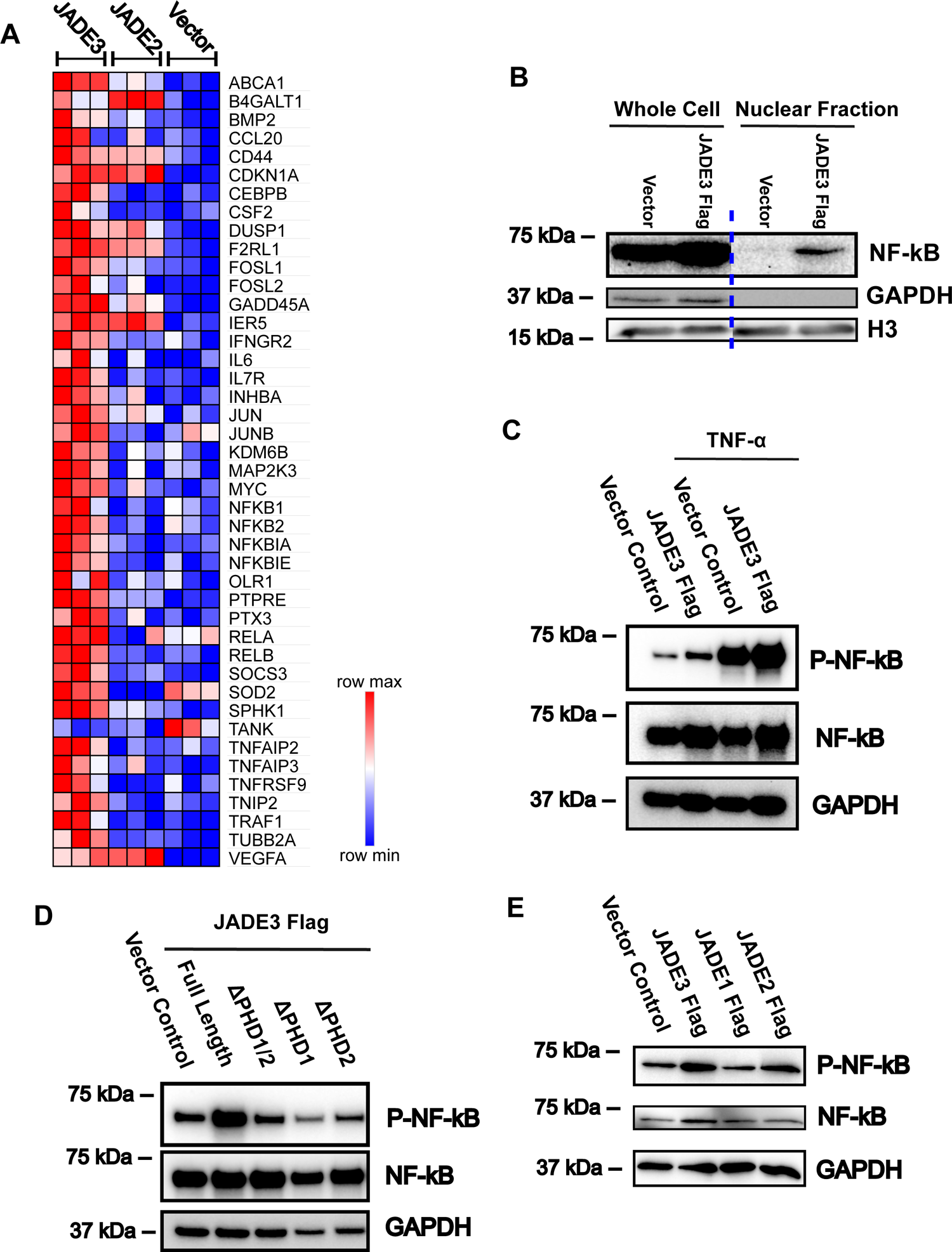
JADE3 activates NF-kB p65. **A.** Heatmap of RNA-sequencing normalized read count for select genes involved in canonical TNF-α/NF-kB signaling pathway. Coloring is done relative to each row with higher normalized read counts in red and lower normalized read counts in blue. Each box is one biological replicate. **B.** Vector control or JADE3 Flag A549 cells were lysed and nuclei were fractionated. Whole cell lysis and nuclear fractions were resolved using SDS-PAGE and probed for NF-kB p65. Image was spliced at blue dashed line from a single blot. Western blot shown is representative of three experiments. **C.** Vector control or JADE3 Flag A549 cells were treated with 20 ng/mL of TNF-α for 15 minutes, then lysed. Whole cell lysis were resolved using SDS-PAGE and probed for phosphorylated NF-kB p65 (Serine 536). Western blot shown is representative of three experiments. **D.** Whole cell lysis from A549 cells expressing either vector control, JADE3 Flag or JADE3 Flag lacking the indicated PHDs were resolved using SDS-PAGE and probed for phosphorylated NF-kB p65 (Serine 536). Western blot shown is representative of three experiments. **E.** Whole cell lysis from A549 cells expressing either a vector control or indicated JADE family members were resolved using SDS-page and probed for phosphorylated NF-kB p65 (Serine 536). Western blot shown is representative of three experiments.

## Discussion

In this study we perform a CRISPRa screen and find a novel antiviral factor JADE3 that activates NF-kB signaling. Our results featuring a well-validated candidate gene list for antiviral activity against IAV provides a resource for those studying influenza biology. Furthermore, our RNA-sequencing dataset provides a foundation for those studying unique and overlapping functions of the Jade family of proteins. Finally, the activation of NF-kB by JADE3 expression presents another mechanism to potentially regulate innate immunity.

Here we conducted a genome-wide CRISPRa screen to identify antiviral factors against IAV. While many CRISPR screens have been conducted to identify host factors involved in viral infection, most have used a loss of function approach. Recently, CRISPR activation has been leveraged to discover novel anti- or pro-viral genes [11, 24, 26, 39–43]. This gain of function approach is well suited for interrogating the immune system as it enables the inclusion of genes that are essential, redundant, or have compensatory mechanisms. As compared to a previous CRISPRa screen done with IAV, we find little overlap between significantly enriched genes [24]. Importantly, our list of 22 potential antiviral genes includes well-known antiviral genes including ISGs and novel genes with unknown antiviral activity against IAV. Many of the enriched genes encode for enzymes that modify glycans (**Figure 1B**). Our results add to the growing body of evidence that glycan modification is an effective antiviral strategy against many viruses [17]. A limitation of our study is that we did not investigate the function of any of these genes, including JADE3 *in vivo*. Thus, further interrogation of the physiological function of these genes during IAV infection *in vivo* is warranted.

The JADE protein family consists of three highly similar proteins that function in the HBO1 complex to direct histone acetylation [36]. This histone acetyltransferase complex is important for bulk H3K14 and H4 acetylation and has largely been investigated in the context of cancer and development [35, 44–46]. The complex consists of multiple subunits including one of either JADE1/2/3 or BRPF1/2/3 using their PHD domains to direct the complex [29]. Furthermore, the switch from BRPF1/2/3 to JADE1/2/3 changes the specificity of the complex from H3K14 to H4 acetylation [29]. To our knowledge comparison of the function of the three JADE proteins has not been done and it has been speculated that they have at least partially redundant function [28]. In this study, we find a unique phenotype for JADE3 expressing cells. JADE3 but not JADE1 or JADE2 significantly inhibit IAV and activate NF-kB signaling. While JADE2 and JADE3 share many differentially expressed genes, the majority of differentially expressed genes are uniquely regulated by JADE2 or JADE3 suggesting non-redundant roles for the JADE proteins. Distinct roles for the JADE proteins is also supported by the phylogenetic analysis which shows distinct orthologs for each of the JADE proteins extending back to jawed vertebrates.

In this study, we identify JADE3 to be a novel regulator of NF-kB activation. NF-kB is an central inflammatory transcription factor that has been heavily studied for its role in inflammatory disease, cancer, and infection [47]. NF-kB functions in both innate and adaptive immunity bridging the transition between the two arms of the immune system and exerting its effects on a variety of cell types including lymphocytes and macrophages [48]. JADE3 expression is elevated in B cells and due to its evolutionary history it is possible the protein also plays a role in the adaptive immune system [49–51]. NF-kB has also been studied for its role in cancer due to its inflammatory, pro-proliferative and anti-apoptotic effects [47]. This is intriguing because JADE proteins are known regulators of the cell cycle and our data shows JADE3 regulating expression of many important pathways in cancer including epithelial-mesenchymal transition, hypoxia, the p53 pathway, apoptosis, glycolysis and KRAS signaling (**Figure 5D and Figure S4**). As a result, the interaction of JADE3, NF-kB, and the other induced pathways in the setting of cancer merits further study.

Owing to its importance in viral infection many viruses either co-opt the NF-kB pathway or inhibit it to support infection [52–55]. Furthermore, overactivation of the pathway has been implicated in the pathogenesis of multiple viruses where damage is often due to over-active inflammatory host damage [56]. JADE3 with its relatively restrained activation of NF-kB and other pro-inflammatory and anti-inflammatory processes may help find the balance in NF-kB activity and merits investigation of its *in vivo* role. This is especially important given that NF-kB has been shown to have both anti-viral and pro-viral roles including with IAV infection [57–59].

Despite its centrality to numerous diseases, many of the genes involved in the immune system are not understood. As one of the natural stimuli for the immune system, viruses can help uncover functionally relevant genes involved in inflammation. Indeed, our study couples the powerful genetic selection viruses exert on host cells with an unbiased genetic screen to assign a functional role to a poorly described human gene with relevance beyond antiviral defense.

## Materials and Methods

### Tissue Culture

293T, A549, HeLa, and MDCK cells were obtained from ATCC and cultured in DMEM supplemented with 5% fetal bovine serum. Cells were routinely screened for mycoplasma contamination. To generate stable cell lines, cells were transduced with lentivirus generated by transfecting lentiviral vectors with psPAX2 and pCMV-VSV-G into 293Ts. After 48 h, supernatants were collected and filtered through a 0.45-μm-pore size filter (Millipore) and added to the respective cells. After 48 hours cells were selected with either puromycin or blasticidin.

To generate A549ΔJADE3 cells, two guide RNAs (sgRNAs: ACGTTCTGTTTATCCGACCC, CCGCTATGACCTAGATGACA) targeting exon 5 of JADE3 were formed using crRNA and tracrRNA from IDT. gRNA were then complexed with Cas9 nuclease and transfected into A549 cells using lipofectamine CRISPRmax (Invitrogen). Cells were then diluted and plated as single cells. Single cell colonies were grown and validated using sanger sequencing for indels resulting in premature stop codons.

### Plasmids

Lenti dCas9-VP64 plasmid (Addgene #61425) was used for CRIPSRa. For validation of the screen guides were cloned into pxPR_502 (Addgene #96923). Human JADE3 (accession no. NP_055550.1), JADE2 (accession no. NP_001276913.1), and JADE1 (accession no. NP_001274369.1) cDNAs corresponding to their protein sequences were cloned with C-terminal 3XFlag tags into pCDH-MCS-T2A-Puro-MSCV (Systems Biosciences, CD522A-1). JADE3 constructs lacking the PHD domains were generated using splicing by overlap extension PCR with PHD1 corresponding to amino acids 200 to 250 and PHD2 corresponding to amino acids 309 to 366, and cloned into pCDH-MCS-T2A-Puro-MSCV. All plasmids were sequence verified prior to use.

### Influenza Virus Assays

A/PR/8/34 was a kind gift of Herbert Virgin (Washington University). mNeon-PR8 has previously been described previously [25]. For PR8 infection, cells were seeded at 100,000 cells per well in a 24 well plate and incubated overnight in culture media. Next day cells were washed with phosphate-buffered saline (PBS) and infected with virus in Virus Production Serum Free Media (VPSFM; Gibco) supplemented with 0.25-0.5 ug/mL TPCK-Trypsin. At specified time points samples were frozen at −80 C.

To determine TCID50 MDCK cells were seeded 25,000 cells per well in a 96 well plate and incubated overnight. Next day cells were washed with PBS and media changed to DMEM+0.2% BSA supplemented with 1.8-2ug/mL TPCK. Cells were infected with serial dilutions of virus samples in replicates and incubated. After CPE was observed (about 3 to 5 days after infection) plates were fixed with a 4% paraformaldehyde solution, stained with crystal violet and scored. TCID50/mL was calculated using Reed-Muench method [60].

### CRISPRa screen

CRISPRa screen was performed similar to what we have previously reported [11]. Briefly, HeLa-dCas9-VP64 cells were transduced with SetA and SetB of the Calabrese CRISPRa library at an MOI of 0.3. Each Calabrese pool was delivered by lentiviral transduction of 1.2 x 10^8^ HeLa cells at an MOI of 0.3. This equates to 3.6 x 10^7^ transduced cells, which is sufficient for the integration of each sgRNA at least 500 independent times. At 2 days post-transduction, puromycin was added to the media, and transduced cells were selected for a total of 7 days. A total of 5 x 10^6^ HeLa CRISPRa cells were seeded in a T-175 tissue culture flask, and each experimental condition was evaluated in duplicate with six independent flasks per replicate (3 x 10^7^ cells per replicate). After seeding cells overnight, media was changed to VPSFM containing 0.5 µg/ml TPCK-Trypsin. Cells either left in TPCK-trypsin containing media or infected with PR8 at an MOI of 1.0 for 48 hours. Two days post-infection, mock-infected cells were harvested for genomic DNA extraction. At this time point, roughly 50% of the PR8-infected cells displayed CPE, and cells were washed twice with PBS, and serum containing media was added. After two days of recovery, cells were again challenged with an additional infection of PR8 at an MOI of 1.0 for 48 hours. Two days after rechallenge, and cells were washed twice with PBS, and serum containing media was added for an addition 3-5 days prior to harvesting. All flasks from a replicate were harvested together. Genomic DNA was isolated from surviving cells using a QIAamp DNA minikit according to the manufacturer’s instructions (Qiagen). Illumina sequencing and STARS analysis were performed as described previously [11].

### Validation

HeLa-dCas9 VP64 cells were transduced with lentiviruses expressing CRISPRa sgRNAs for the indicated genes (**Figure S6**). Transcriptional activation was confirmed via qPCR for the corresponding gene. Briefly, RNA was extracted using a Direct-zol kit (Zymo Research) and cDNA prepared using a High-Capacity cDNA Reverse Transcription Kit (Thermo). qPCR was performed using primers and probes from IDT (**Figure S7**). Expression was normalized relative to the mRNA expression of actin. Cells were then seeded 100,000 cells per well in a 24 well plate and incubated overnight. Cells were then washed and infected with PR8 mNeon virus in VPSFM supplemented with 0.5 ug/mL of TPCK-Trypsin. Cells were then incubated in the Incucyte live cell analysis instrument (Sartorius) and green fluorescence and phase images were captured every 3 hours for 48 hours total. Fluorescent (infected) cells were quantified using the Incucyte software.

### RNA-sequencing

Cells were seeded 200,000 cells per well in a 12 well plate and incubated overnight. Next day cells were washed and media changed to VPSFM supplemented with 0.5 ug/mL TPCK-Trypsin. After 4 hours RNA was isolated (Direct-zol). mRNA sequencing was performed by Novogene using an Ilumina platform and polyA capture. Reads were aligned using HISAT2 and differential gene quantification done using DESeq2 [61, 62]. Principal component analysis was performed using DESeq2. Gene set enrichment analysis was done using the Molecular Signatures Database hallmark gene sets [63, 64]. Matrix analysis was done using Morpheus software (https://software.broadinstitute.org/morpheus).

### Western blot

Cells were seeded and incubated overnight in culture media. Cells were lysed in laemlli buffer supplemented with beta-mercaptoethanol and HALT protease and phosphatase inhibitor (PPI) cocktail (Sigma). Isolation of nuclei from whole cell lysis adapted a standard centrifugation based protocol [65]. Briefly, cells were lysed in 0.1% Igepal in PBS supplemented with PPI cocktail. Nuclear fractions were sedimented by centrifugation. Whole cell lysates and nuclear fraction were resolved using SDS page. Western blot was done using a standard protocol and images processed using ImageJ. For phospho-NF-kB experiments cells were serum starved for 4-8 hours before lysing. Recombinant human TNF-α (R&D systems 210-TA) treatment was done at 20 ng/mL for 15 minutes before lysing cells. Antibodies used include: Rabbit anti-JADE3 (Abcam #129495) used at 1:1000, Rabbit anti-NF-kB p65 (Cell Signaling #8242) used at 1:1000, Rabbit anti-phospho-NF-kB p65 Ser536 (Cell Signaling #3033) used at 1:1000, Rabbit anti-H3 (Cell Signaling), anti-Flag HRP (Sigma), anti-rabbit IGG HRP (Sigma), anti-GAPDH HRP (Sigma)

### Phylogenetic and sequence analysis

Ortholog gene groups for each of the JADE proteins were identified using NCBI orthologs (https://www.ncbi.nlm.nih.gov/kis/info/how-are-orthologs-calculated/). For Lamprey reciprocal blast hits were performed to find potential orthologs. Amino acid sequences were retrieved from NCBI (accession numbers in **Figure S8**) and used for multiple sequence alignments using MAFFT [66, 67]. Maximum likelihood trees were inferred by PhyML using 100 bootstrap replicates [68, 69]. Drosophila Melanogaster Rhinoceros protein was used as an outgroup to root tree. Protein sequence alignment of human JADE proteins was done using NCBI’s COBALT [70].

## Data Availability

Raw mapped read counts and STARS analysis from the CRISPRa screen are available in the supplemental tables. Normalized read counts and differentially expressed genes for all RNA sequencing from this study are also in the supplemental tables. All raw RNA sequencing data is deposited in the NCBI sequence read archive (SRA) under the BioProject ID: PRJNA1022046.

## Acknowledgements

We would like to thank Dr. Peter Palese (Icahn School of Medicine at Mount Sinai) for his generous contributions of IAV strains and resources. We also would like to thank Dustin Hancks, John Schoggins, Julie Pfeiffer, and all members of the Orchard Lab for helpful discussions. M.M. was supported by the UT Southwestern Molecular Microbiology Training Grant (T32 AI007520). R.C.O. was supported by NIH grant R35GM142684.

## Author Contributions

M.M. designed the project, performed experiments, and helped draft the paper. A.E. and J.G.D. performed experiments and analyzed data. P.P., N.S.H., C.W.B., provided critical reagents and expertise for all influenza A viral experiments. R.C.O. conceptualized the project, provided supervision, and helped write the paper. All authors read and edited the manuscript.

## SUPPLEMENTAL INFORMATION APPENDIX

### Supplemental Tables

**Table S1:** STARS analysis of genome-wide CRISPRa screen in HeLa cells for IAV

**Table S2:** Raw Mapped Read counts from the mock and PR8 infected pools of the genome-wide CRISPRa Screen

**Table S3:** Results from RNA sequencing A. Differentially expressed genes from RNA-sequencing of JADE3 Flag vs vector control expressing cells B. Differentially expressed genes from RNA-sequencing of JADE2 Flag vs vector control expressing cells C. Differentially expressed genes from RNA-sequencing of JADE3 Flag vs JADE2 Flag expressing cells D. Normalized read counts as fragments per kilobase of transcript per million mapped reads (FPKM) from RNA-sequencing of JADE3 Flag, JADE2 Flag and vector control expressing cells

**Figure S1:**
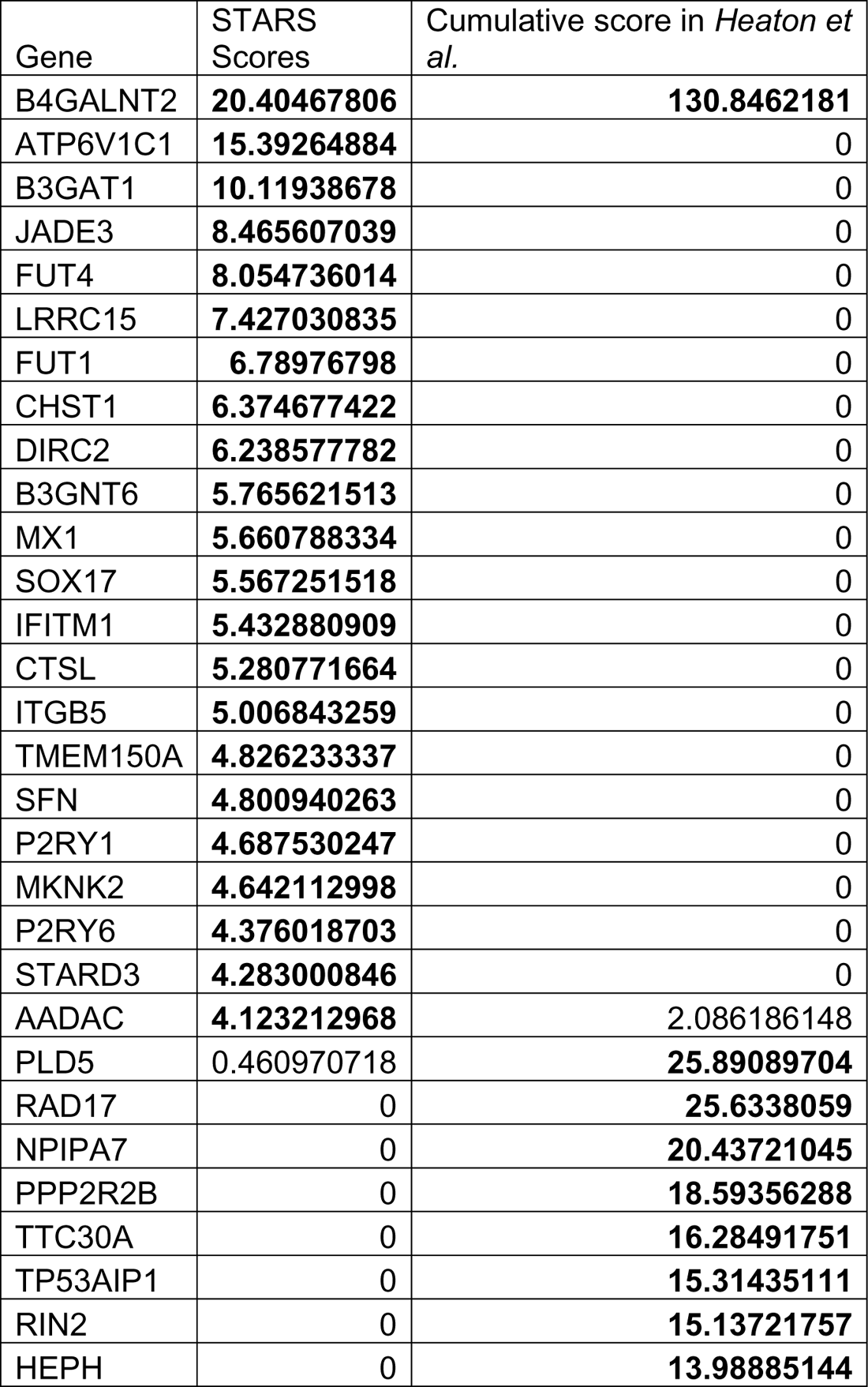
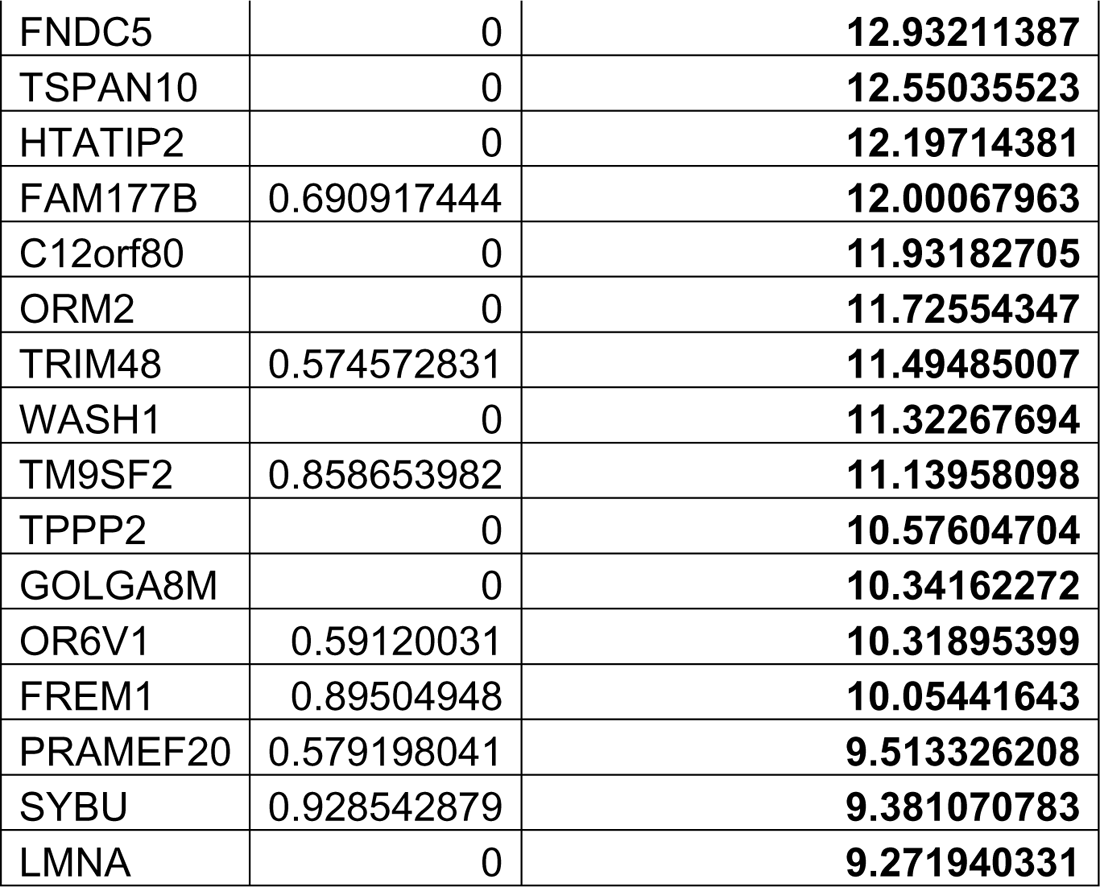
Comparison of significantly enriched genes between IAV CRISPRa screens. STARS score (current study) and Cumulative score (*Heaton et al.*) are plotted for all genes that reached statistical significance in either study. Bolded values indicate statistical significance as defined by each study. A score of 0 denotes unscored.

**Figure S2:**
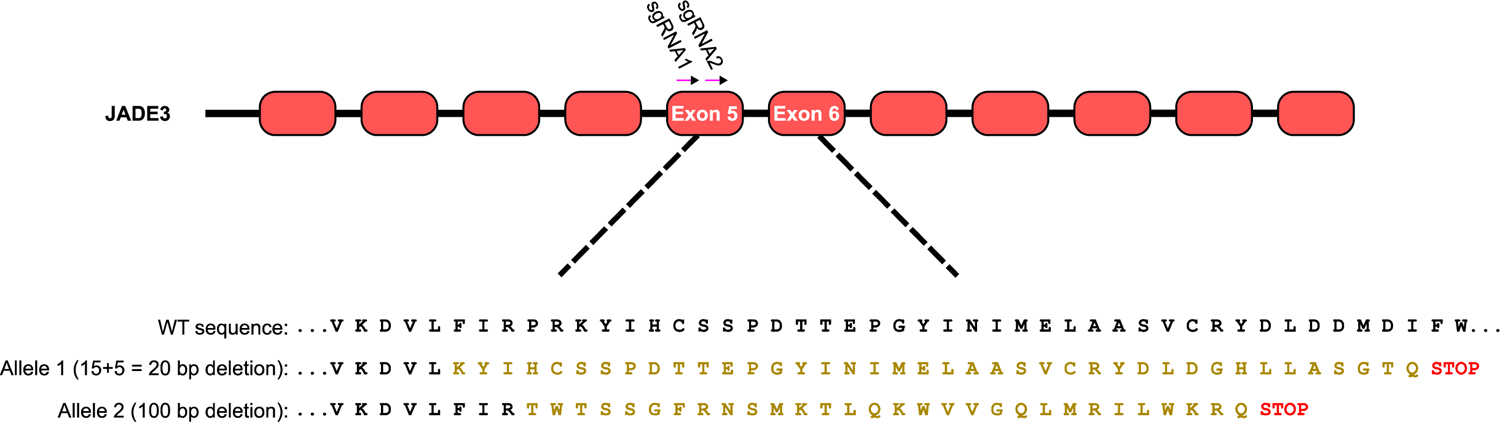
Two gRNA (pink) targeting exon 5 of JADE3 and Cas9 were transfected into A549 cells. Single cells colonies were sequenced. Wild type amino acid sequence is shown as well as the amino acid sequence corresponding to the sequencing of allele 1 and 2 of A549ΔJADE3 cells resulting in premature stop codons.

**Figure S3:**
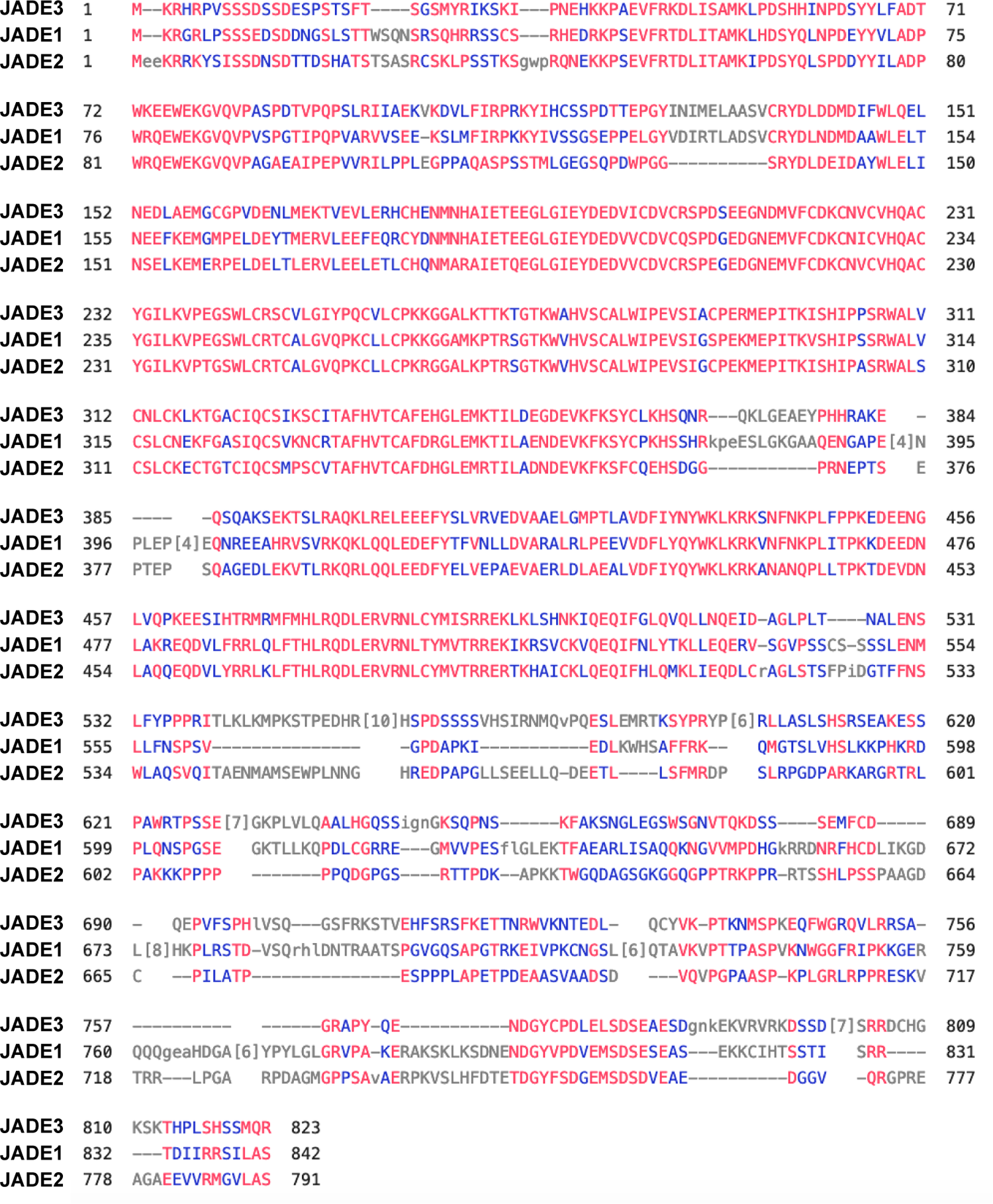
COBALT sequence alignment of Human JADE3 (accession no. NP_055550.1), JADE2 (accession no. NP_001276913.1), and JADE1 (accession no. NP_001274369.1). Unaligned columns are compressed into the bracket form denoting the number of unaligned residues. Coloring is based on a conservation threshold of 3 bits, amino acids in red are highly conserved while those in blue are less conserved.

**Figure S4:**
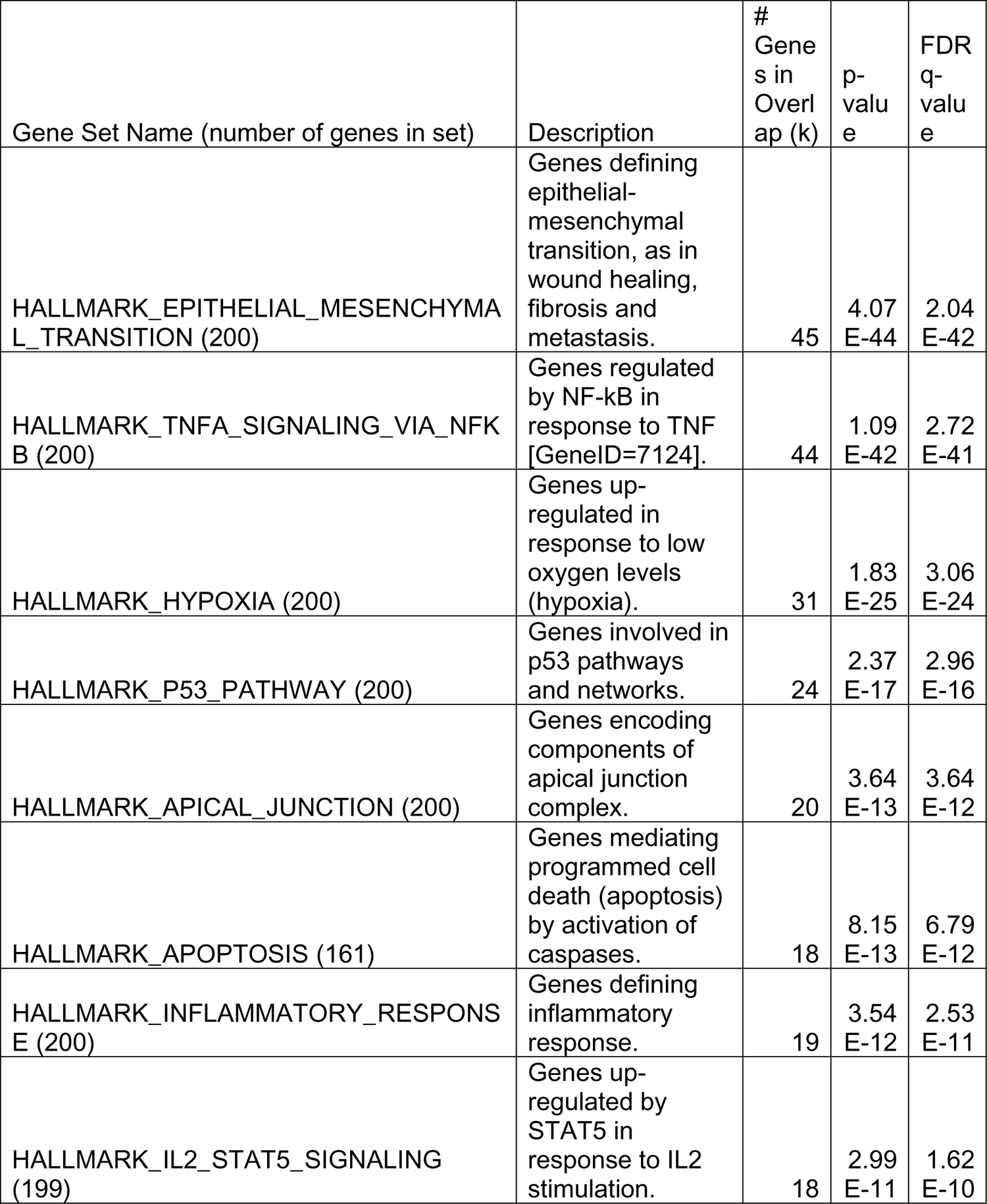
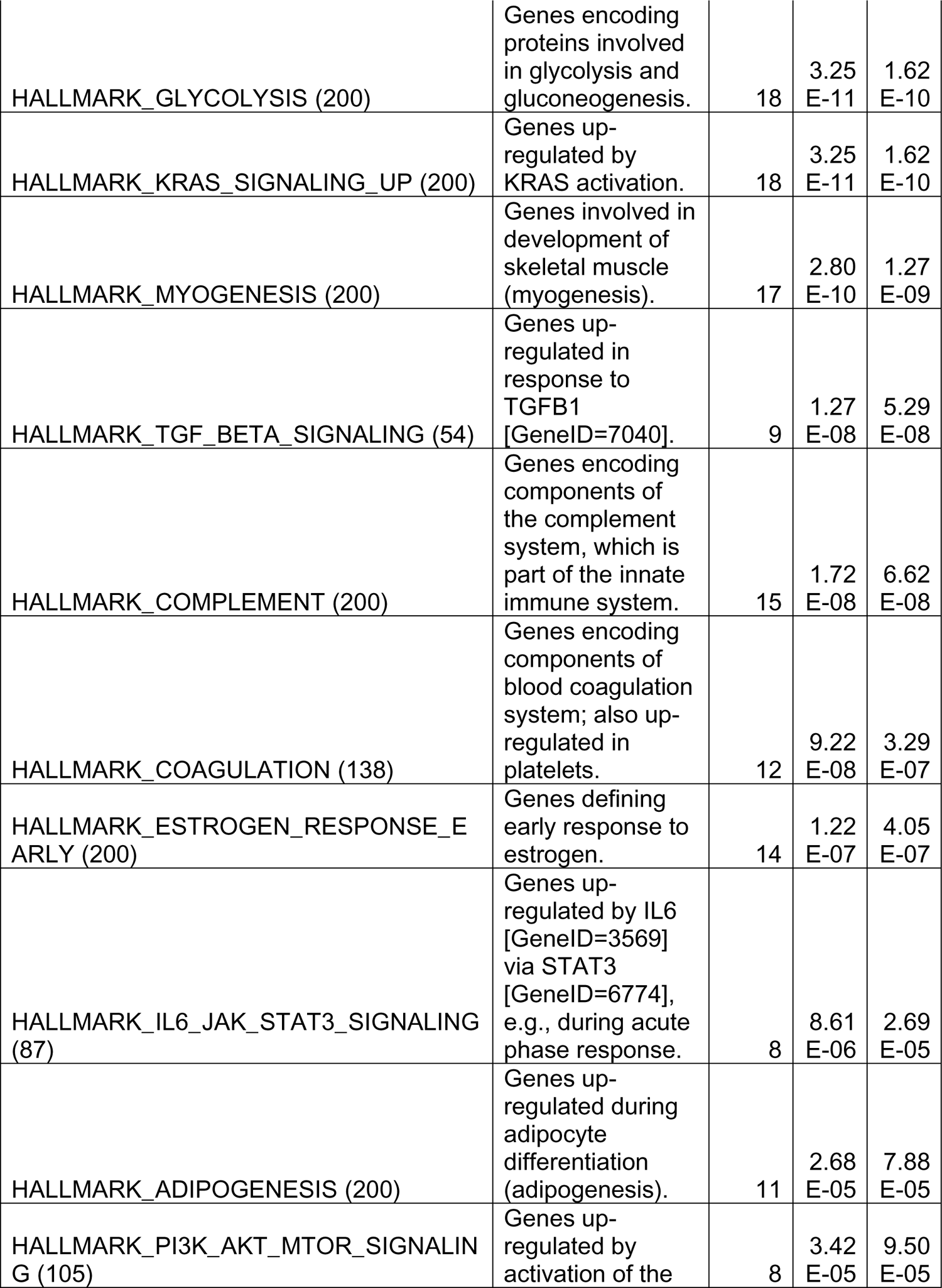
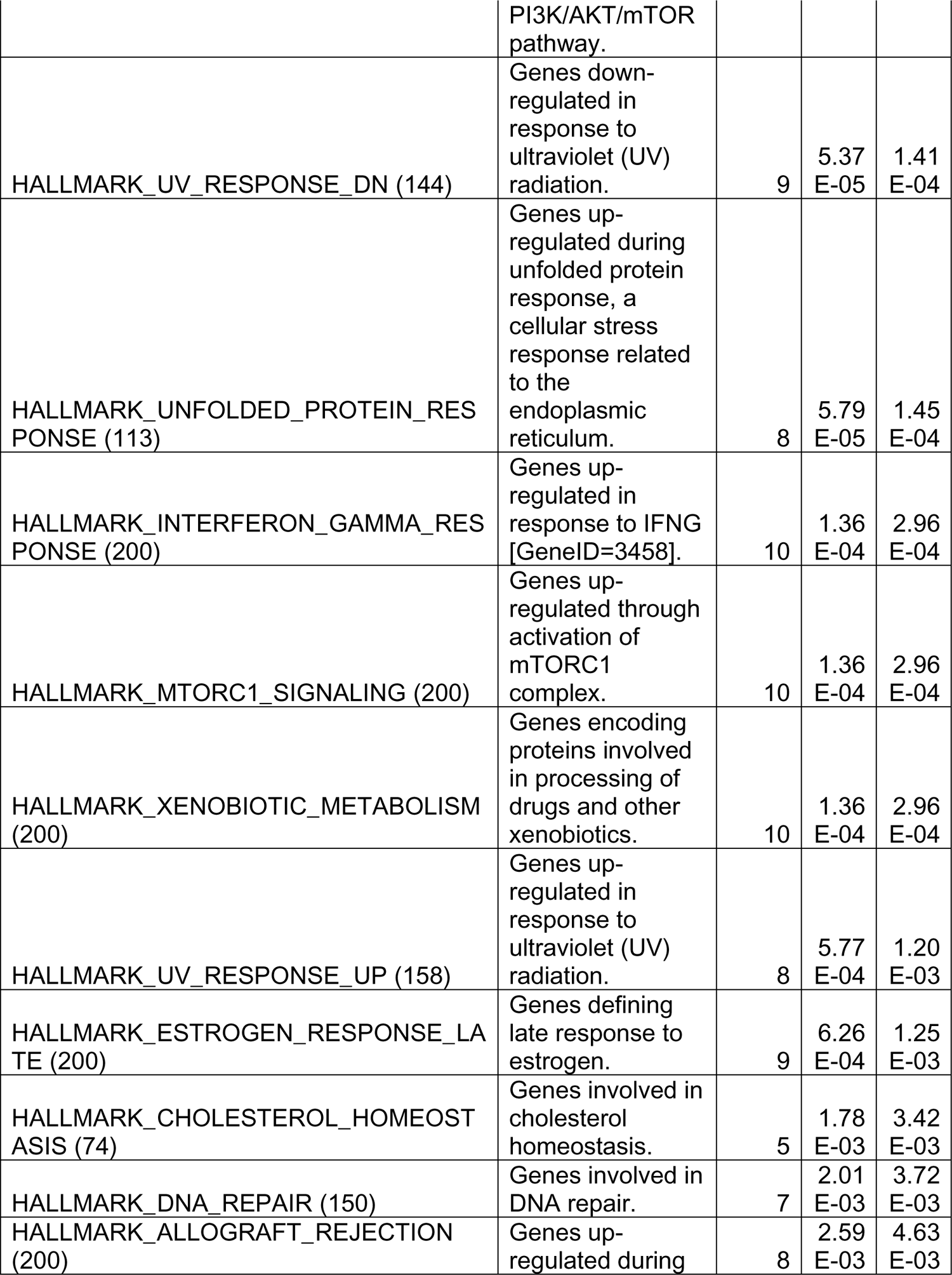
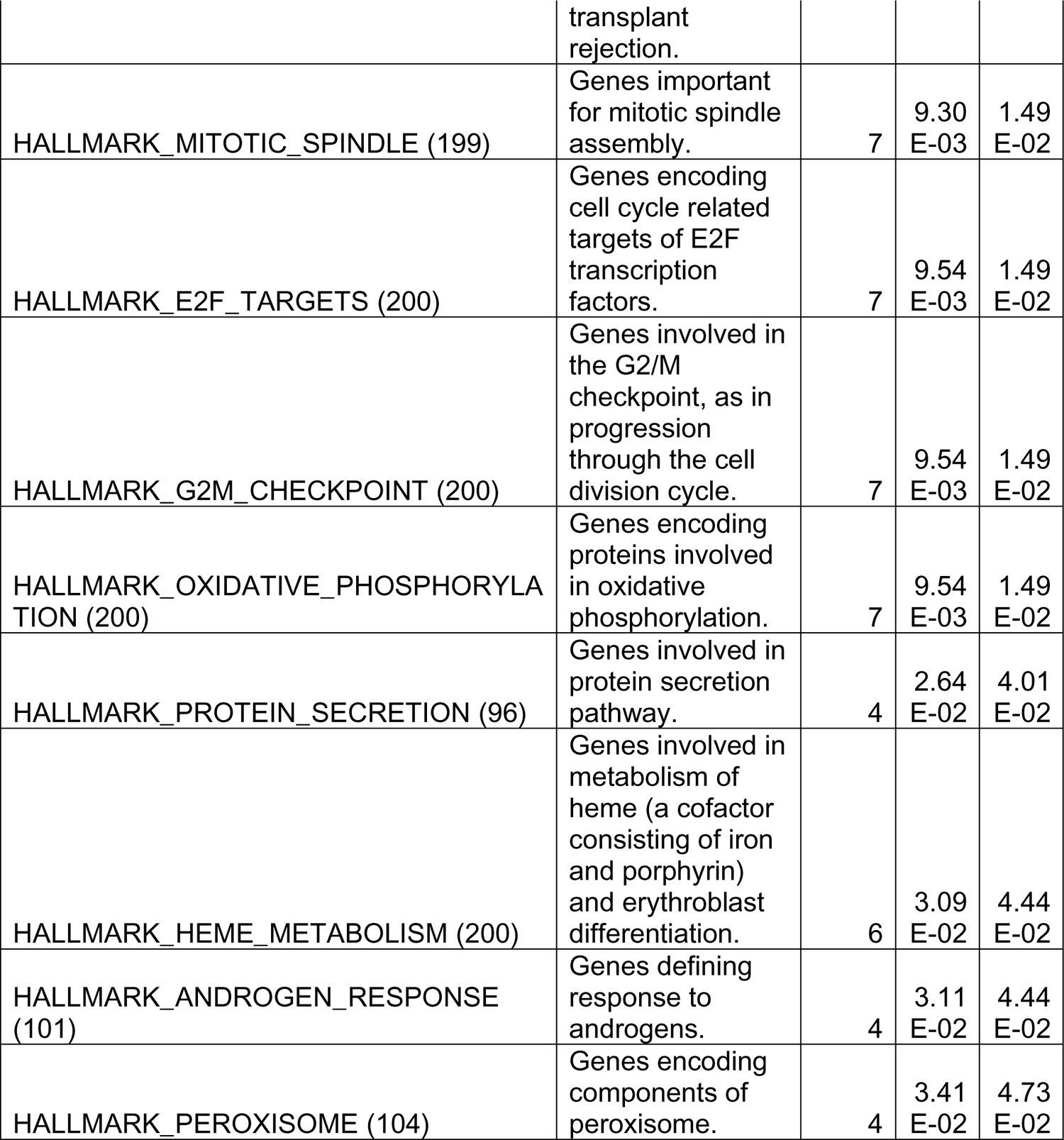
Expanded gene set enrichment analysis of the 500 most significantly upregulated genes in JADE3 Flag vs vector control A549 cells.

**Figure S5:**
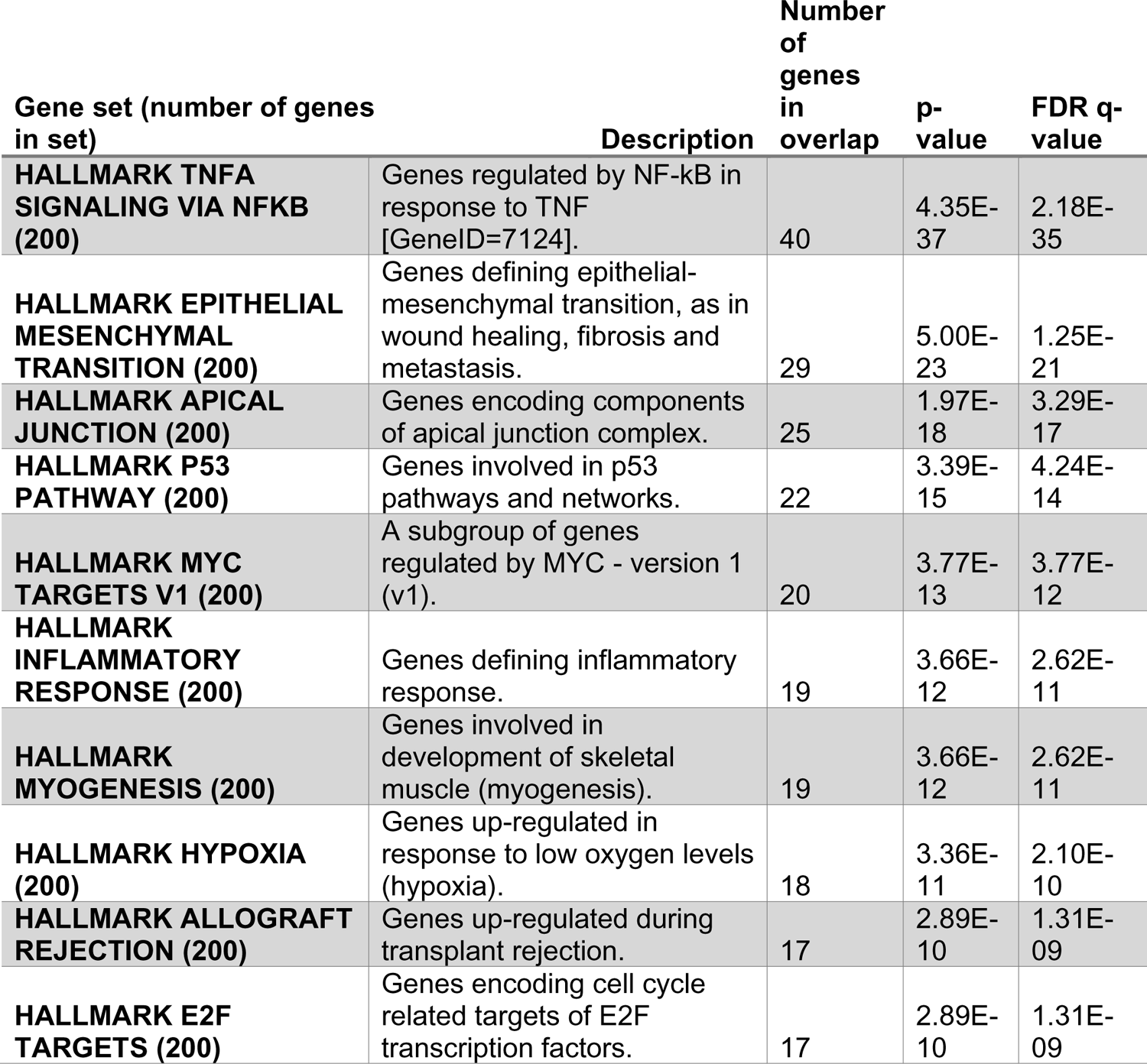
Gene set enrichment analysis of the 500 most significantly upregulated genes in JADE3 Flag vs JADE2 Flag A549 cells. The 10 most significantly enriched gene sets are shown

**Figure S6:**
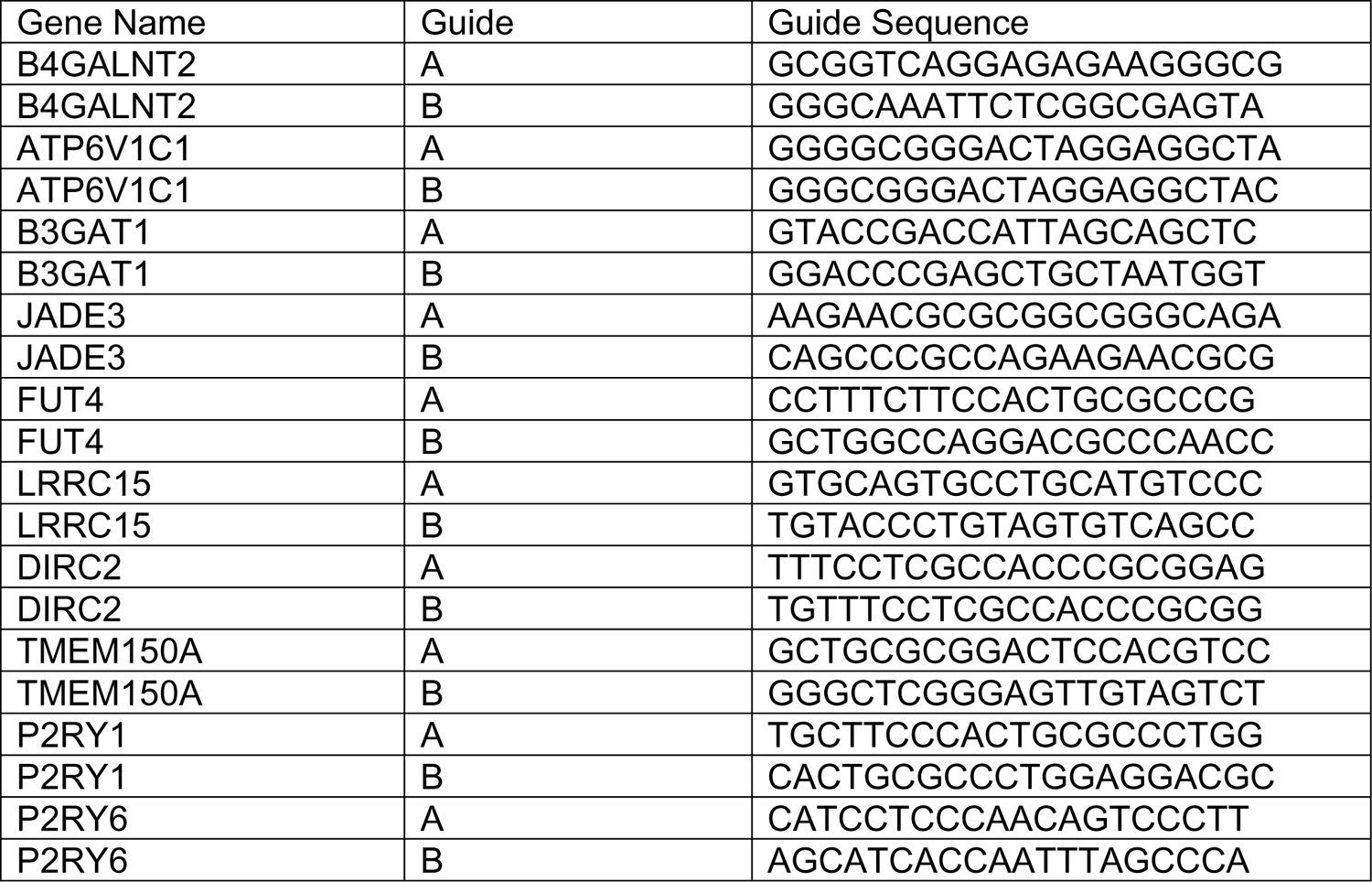
sgRNAs used in this study for CRISPRa validation.

**Figure S7:**
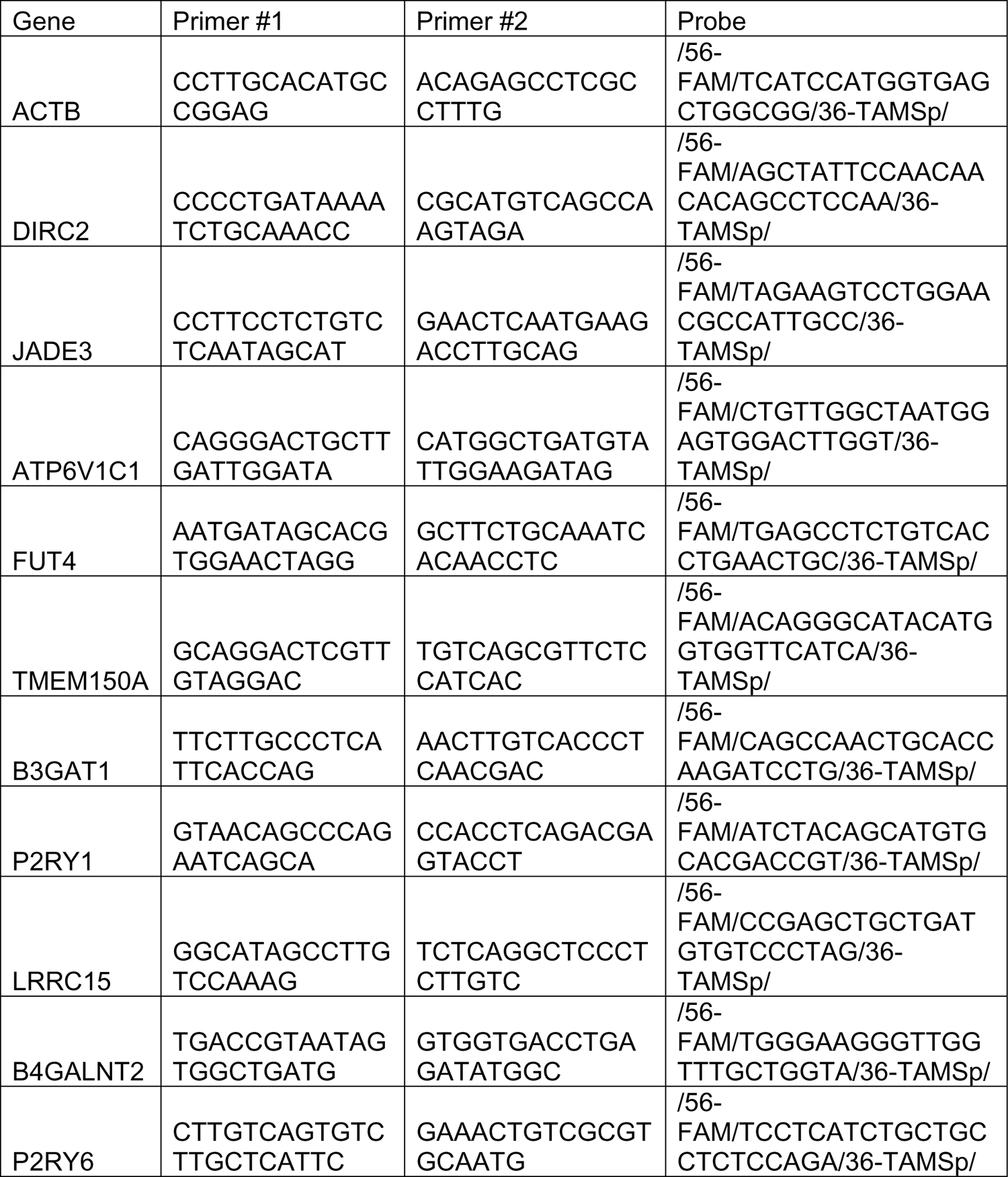
Primer and probe sequences for qPCR

**Figure S8:**
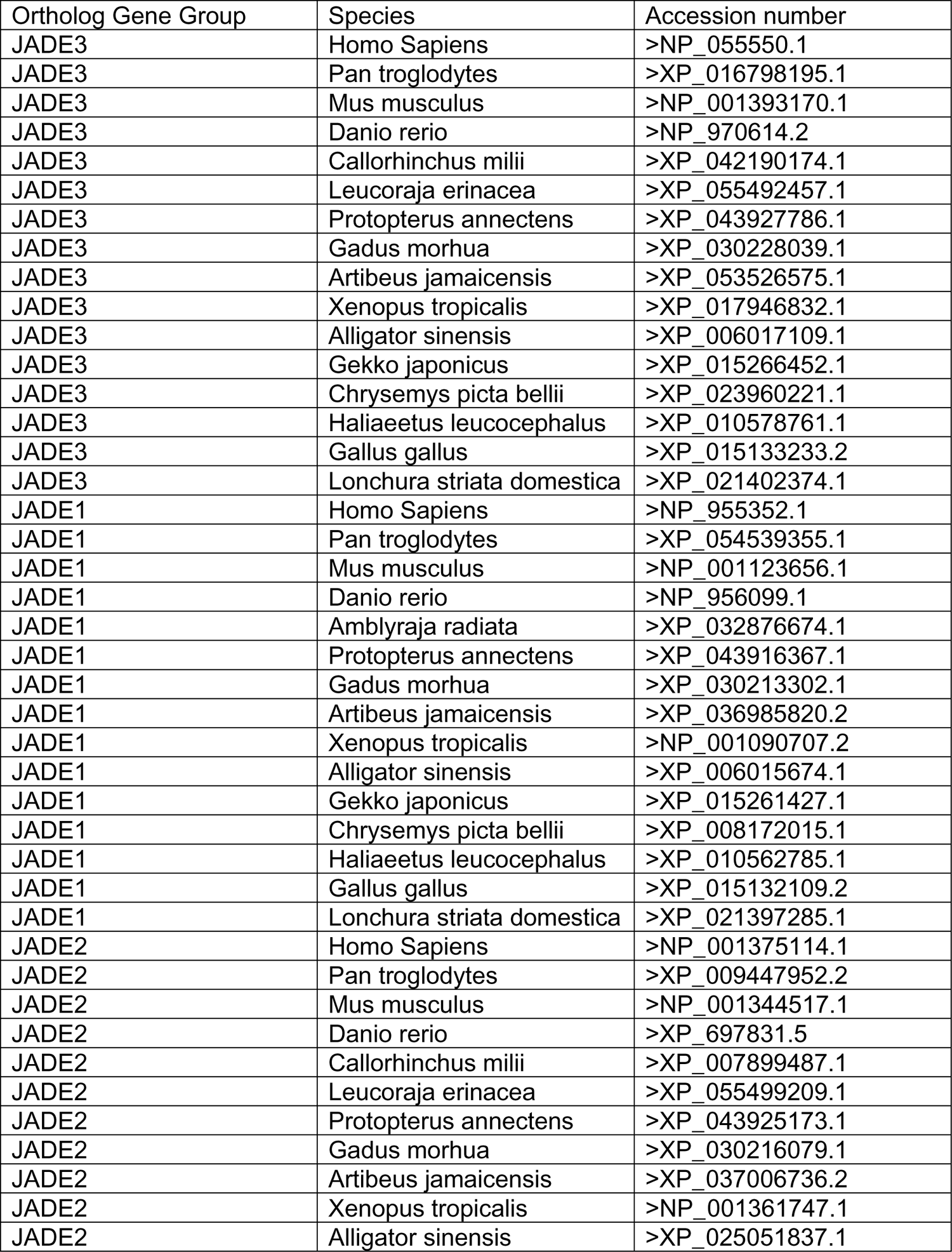
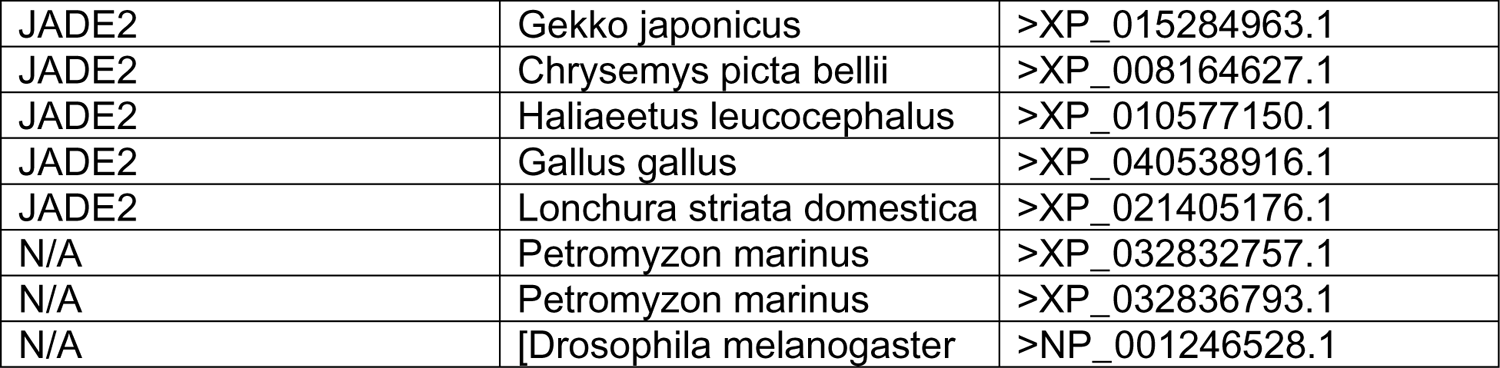
Accession numbers for sequences used for phylogenetic analysis

